# Pupil-linked phasic arousal relates to the reduction of response inhibition: inferences from a simultaneous pupillometry and EEG-fMRI study

**DOI:** 10.1101/2022.08.22.504728

**Authors:** Linbi Hong, Hengda He, Paul Sajda

## Abstract

Attention reorienting is a critical cognitive function which drives how we respond to novel and unexpected stimuli. In recent years, arousal has been linked to attention reorienting. The timing and spatial organization of the interactions between the arousal and reorienting systems, however, remain only partially revealed. Here, we investigate the dynamics between the two systems through simultaneous recordings of pupillometry, EEG, and fMRI of healthy human subjects while they performed an auditory target detection task. We used pupil diameter and activity in the noradrenergic locus coeruleus to infer arousal, and found these measures linked to distinct cortical activity at various temporal stages of the reorienting response. Specifically, our results provide the first demonstration in humans of how phasic pupil-linked arousal relates to the reduction of response inhibition, an inference which otherwise would remain hidden without the help of simultaneous multi-modal acquisitions.

## Introduction

Attention reorienting is one of the most fundamental and critical cognitive functions for both animals and humans. The ability to redirect attention to certain novel and unexpected stimuli enables us to adapt to and interact with the environment in a highly efficient manner ^1-3^. Successful reorienting allows us to timely avoid danger and reap rewards, whereas impaired reorienting can impede a spectrum of daily activities (for instance in populations with autism or depression ^4-7^). Previous work indicates that arousal modulates the reorienting response through a coordinated action of both cortical and subcortical systems ^1,8,9^. The exact spatiotemporal dynamics of such modulation, however, remains less clear. This work describes a multi-modal approach which enables comprehensive examination and interpretation of dynamics between the arousal and reorienting systems. Techniques and inferences developed from this study can therefore yield insights for both nonclinical and clinical research.

One of the major challenges in investigating the dynamics between arousal and attention reorienting originates from the complex physiological basis of arousal. As an index for the level of wakefulness, arousal is regulated by a number of subcortical brainstem structures ^8,9^. Fluctuations in cortical arousal are thus enabled via the widespread release of neurotransmitters (such as acetylcholine and norepinephrine) from the brainstem to the cortex ^8-12^. In in-vivo animal studies, dynamics of arousal can be monitored with high fidelity via direct recordings at brainstem nuclei such as the noradrenergic locus coeruleus (LC) and cholinergic basal forebrain (BF) ^8,11,13^. In noninvasive human studies, however, acquiring high quality recordings from brainstem structures is particularly challenging. This is mainly due to the difficulties in acquisition, and potential contaminations on acquired signal from physiological noise ^14-17^. This problem has in part precluded direct investigation of arousal in human studies, and motivated researchers to look for indirect yet robust markers indicative of internal arousal state.

Pupil diameter under constant luminance emerged as one of the widely used markers to index arousal ^8,18-21^. In recent human studies where high quality recordings from brainstem structures were successfully acquired, a tight link has also been observed between changes in pupil diameter and responses in brainstem nuclei such as the LC and BF ^12,22^. Along with previous evidence from animal studies reporting synchrony between pupil diameter and the activity of neuromodulatory nuclei ^8,13,23^, these findings demonstrate the reliability of pupil as a proxy for the internal arousal state.

Many groups have capitalized on the convenience of pupillometry and used it concurrently with scalp electroencephalography (EEG) or functional magnetic resonance imaging (fMRI) to infer the relationship between pupil-linked arousal and attention relevant neural processes ^12,18,22,24,25^. Simultaneous pupil-fMRI studies have identified spatial correlates of pupil-linked arousal such as the frontal eye field, anterior cingulate cortex, and visual cortex ^22,26,27^. Meanwhile, simultaneous pupil-EEG studies report that tonic pupil-linked arousal correlates with P3, an event related potential (ERP) indicative of attention reorienting towards less frequent target stimuli ^18,24,25^.

While recent work has furthered our understanding of the dynamic interactions between the arousal and reorienting systems, measures used in previous studies limit the inferences that can be drawn. For instance, in pupil-fMRI studies ^12,22^, both pupil diameter and blood oxygenation level dependent (BOLD) signal fluctuates on timescales of seconds. It is therefore difficult for these measures to capture millisecond-level reorienting relevant processes, and to pinpoint the exact timing when interactions between the arousal and reorienting systems take place. Meanwhile, in pupil-EEG studies ^18,24,25^, a large number of analyses investigate the dynamics between systems by correlating amplitudes of ERPs to pupil diameter. ERP amplitudes are generally computed by averaging the maximal deviation of evoked responses across trials (e.g., amplitudes of the P3 component are generally extracted from 250 to 500 ms after stimulus onset). While ensuring the signal-to-noise ratio of EEG, this approach dilutes the millisecond resolution of EEG. It also masks trial-by-trial variability, which may be important for investigating dynamics that are independent of the stimulus. Moreover, in pupil-EEG studies where temporal specificity and trial-by-trial variability of EEG are indeed preserved, the suboptimal spatial resolution of EEG makes it difficult to identify precise locations of interactions.

To address these limitations, we used a simultaneous acquisition to capture physiological and neural activity at both high temporal and spatial resolutions. Specifically, we simultaneously measured pupillometry, EEG, and fMRI of healthy human subjects while they performed an auditory oddball task inside the MR scanner. The auditory oddball paradigm was used to minimize ocular confounds and facilitate fluctuations of internal arousal. We operationalized arousal in terms of pupil diameter and functional BOLD activity of the LC, and investigated relationships between arousal and reorienting relevant systems. First, we used temporally specific EEG measures to inform fMRI analysis. This allowed us to identify spatial correlates of reorienting relevant cortical activity with millisecond resolution. Second, we defined results from the EEG-fMRI analysis as cortical regions of interest (ROI), and examined relationships between ROI-extracted BOLD signal and both baseline pupil diameter (BPD) and task-evoked pupil response (TPR). With the baseline and evoked pupillary measures indicative of tonic (i.e., slow and spontaneous) and phasic (i.e., fast and task-evoked) arousal respectively, this analysis assessed the time and prevalence of relationships between pupil-linked arousal and reorienting relevant cortical activity with high spatiotemporal precision. Next, we evaluated the relationship between LC-linked arousal and reorienting with an analysis analogous to the EEG-informed fMRI analysis. Specifically, in this analysis, temporally specific EEG measures were correlated to task-evoked LC responses (whereas in the EEG-informed fMRI analysis, the EEG measures were correlated to BOLD responses in the cortex). Lastly, we evaluated the extent to which pupil reflects LC activity by quantifying task-evoked LC response with pupillary measures.

In summary, we first used ROIs resulted from the EEG-fMRI analysis to parse the timing of cortical activity and lay out the cortical dynamics. We then correlated activity at these ROIs to pupillary measures and LC responses to link the cortical dynamics to arousal. With this approach, we found that phasic and tonic pupil-linked arousal interact with the reorienting response in a coordinated fashion. Specifically, our findings revealed that ROI-extracted BOLD signal at 225 ms and 275 ms poststimulus were linked to TPR, whilst ROI-extracted BOLD signal at 275 ms and 425 ms poststimulus were linked to BPD. On the other hand, LC activity was not linked to cortical activity located at those exact windows, but at windows in close temporal vicinity (i.e., at 250 and 300 ms). Meanwhile, we found LC activity to be correlated with BPD, confirming the tight link between pupil, LC and cortical arousal. Taken together, these findings enabled inferences on the potential role of phasic pupil-linked arousal in reducing response inhibition, and on the role of tonic pupil-linked arousal in favoring exploration over exploitation. It also allowed us to tease apart the common and unique cortical activity captured by both pupil- and LC-linked arousal, and to provide one of the first lines of evidence on inferring the specific roles played by task-evoked and spontaneous pupil-linked arousal in modulating cortical activity at different temporal stages of attention reorienting.

## Results

We collected simultaneous pupillometry, EEG, and fMRI data from 19 participants while they performed an auditory oddball task inside the MR scanner (Fig. 1a and 1b). Subjects were instructed to respond with a right hand button press once they heard the target (oddball) sound, and to maintain a stable fixation to a crosshair on the screen throughout the experiment. Overall, subjects completed the task with high accuracy (99.4% ± 0.1%), and responded to the target stimuli with an average response time (RT) of 414.7 ± 69 ms.

**Figure 1.**
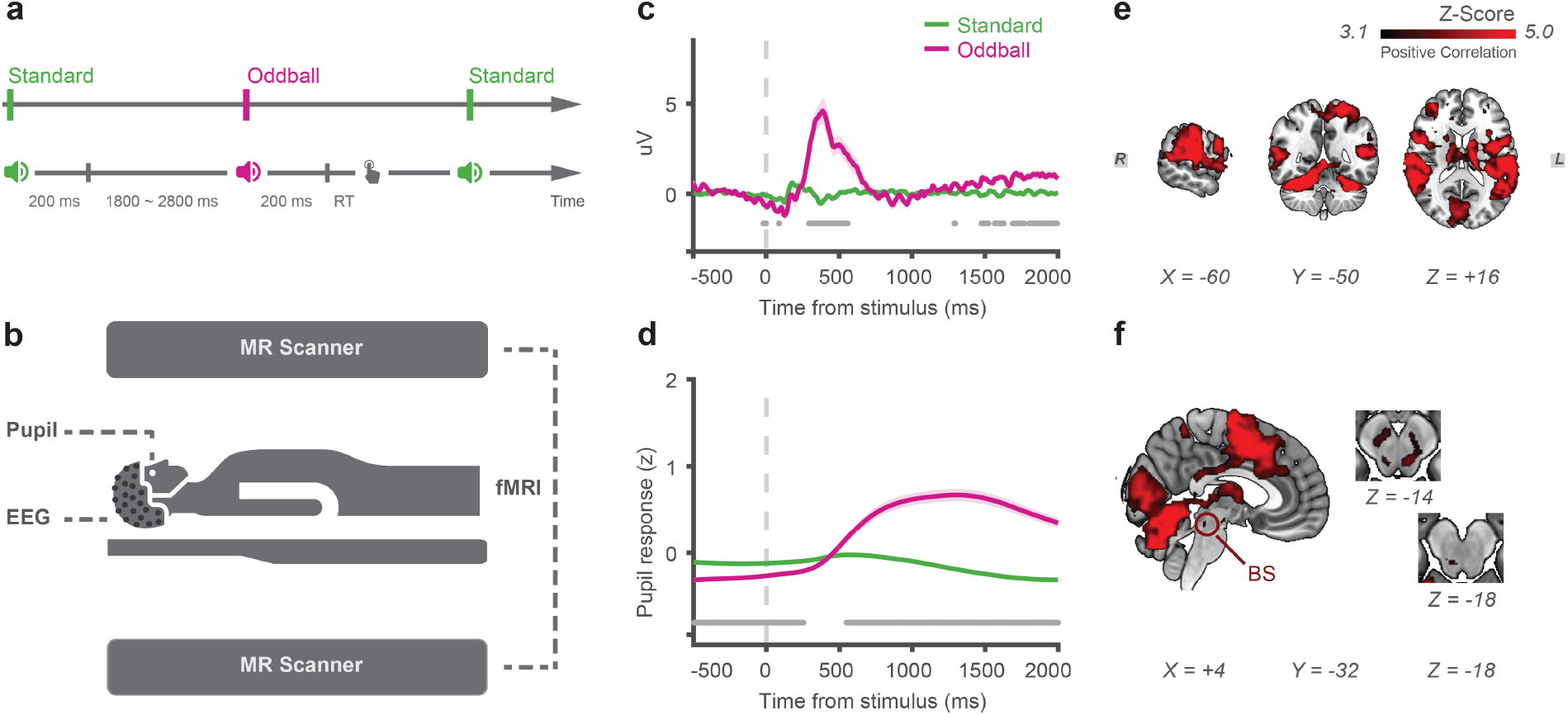
Simultaneous acquisition of pupillometry, EEG, and fMRI with an auditory oddball paradigm. (**a**) Experimental design. Subjects were instructed to maintain fixation to a fixation target on the screen, and press a button once they heard the oddball (target) sound (20% of occurrence rate). Each trial was composed of a 200 ms stimulus presentation, and a 1800 to 2800 ms period for subjects to respond. (**b**) Acquisition environment. Multi-modal data were acquired concurrently inside the scanner. (**c**) Time course of trial-averaged ERP at Pz. Stimulus onset is at time 0, with dark gray line at the bottom indicating significant difference (*P* < 0.001) between ERPs of different stimulus types. (**d**) Time course of trial-averaged z-scored pupillary response. (**e**) Traditional fMRI analysis results showing greater BOLD response to target than standard stimuli in the cortical and (**f**) brainstem regions. X, Y, Z are Montreal Neurological Institute (MNI) coordinates. BS, brainstem. All panels: group average (*N* = 19); shading, s.e.m.; statistics, one-sample t-test (panels **c** and **d**) or Gaussian random field theory (panels **e** and **f**).

**Figure S1. Related to Figure 1.**
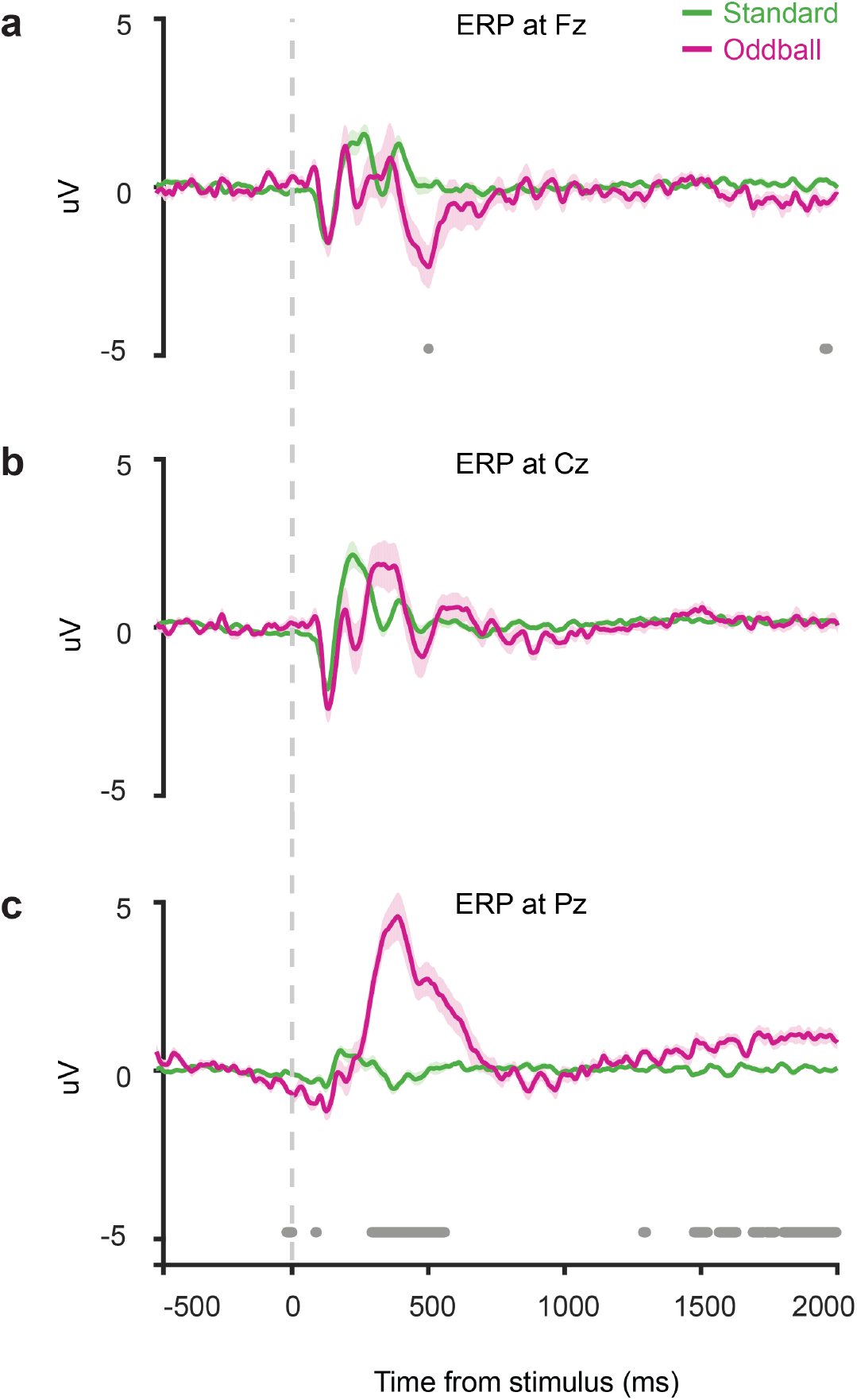
Trial-averaged evoked potentials at Fz (**a**), Cz (**b**), and Pz (**c**) from top to bottom. Stimulus onset is at time 0, with dark gray line at the bottom indicating significant difference (*P* < 0.001) between ERPs of different stimulus types. All panels: group average (*N* = 19); shading, s.e.m.; statistics, one-sample t-test.

**Figure S2. Related to Figure 1.**
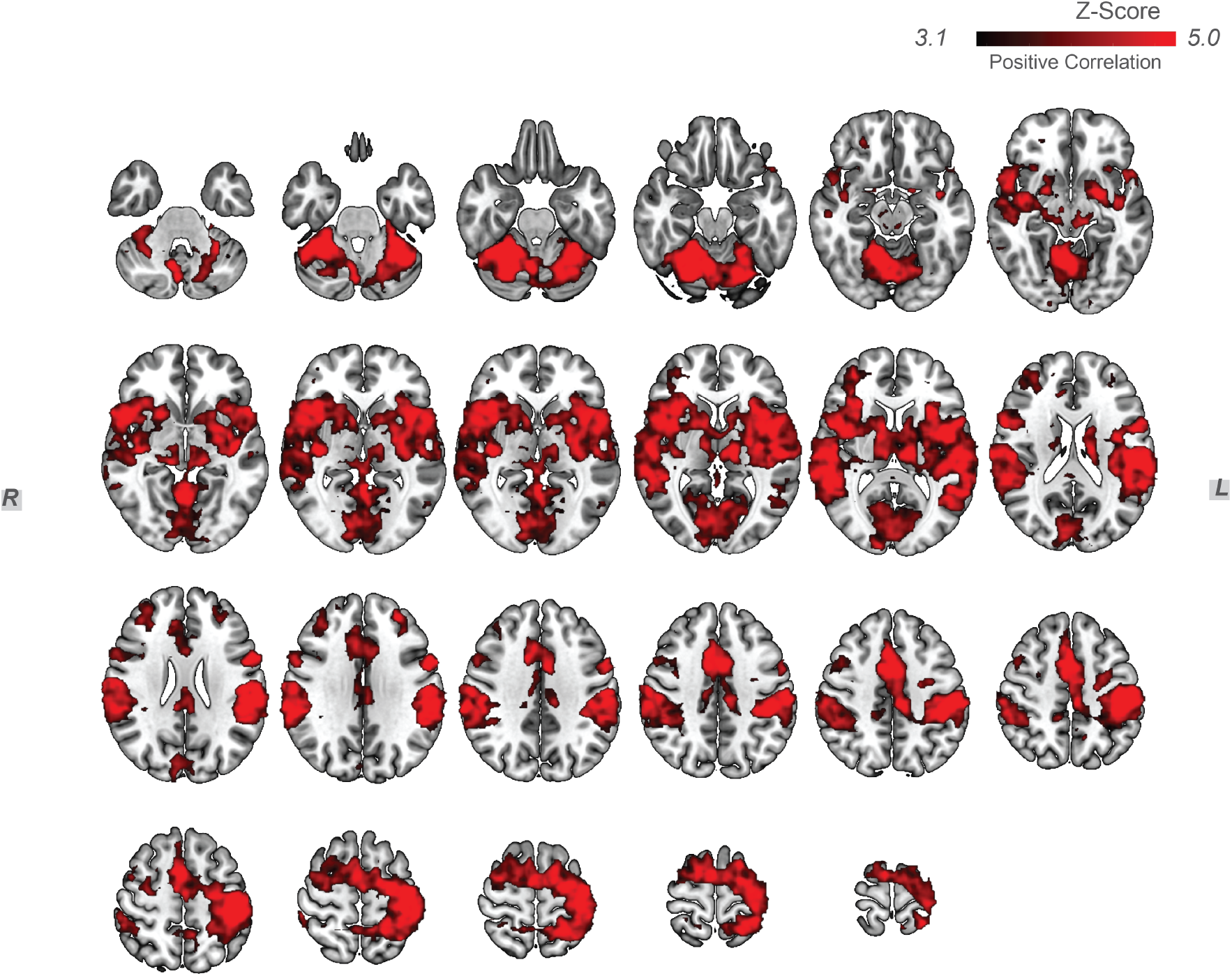
Traditional fMRI analysis results for the oddball (target) > standard contrast (i.e., activation maps showing greater BOLD response to oddball than standard stimuli). All results were multiple comparisons corrected with *z* > 3.1 and *P* < 0.05.

### Single modality results were consistent with previous findings

Despite the technical challenges in data acquisition and the high-amplitude artifacts introduced to EEG by the MR scanner, trial-averaged ERPs showed typical patterns as observed in previous studies ^19,28-31^. The typical sequence of ERPs elicited by an auditory oddball paradigm were all present: N1, P2, N2, and P3, as shown at three midline sites (Fz, Cz, and Pz) in Fig. S1.

Similarly, consistent with previous findings ^18,19^, evoked changes in pupil diameter did not take place until a few hundred milliseconds into the trial (Fig. 1d). Meanwhile, traditional fMRI analysis revealed a distributed network of areas showing stronger BOLD responses for target versus standard stimuli (Fig. 1e). Activations in cortical regions involved in sensory processing (Heschl’s gyrus), decision making (prefrontal gyrus, cingulate cortex) and motor response (precentral gyrus) were all present (Fig. S2), as reported in previous studies ^32,33^. Interestingly, a subset of brainstem structures also showed greater BOLD for target versus standard stimuli (Fig. 1f). This observation supports recent findings on the role of subcortical neuromodulatory systems in attention modulation and reorienting ^12,13,34^.

### EEG-informed fMRI analysis revealed cascade of cortical activity linked to single trial variability in the reorienting response

To identify temporally specific neuronal components associated with different task conditions, we performed a single-trial analysis on the EEG signal ^35^. Specifically, for each subject, we trained a classifier to discriminate evoked responses of targets from standards at a number of poststimulus windows within the time span of the trial. At each predefined temporal window, the classifier estimated a set of spatial weights that achieves maximal discrimination between target versus standard conditions. These weights were then applied to single-trial EEG data to capture trial-by-trial amplitudes of EEG discriminating components (Fig. S3). Performance of the classifier was estimated with a receiver operating characteristic curve (ROC) using a leave-one-out cross validation procedure ^35,36^, and was significant from 125 to 950 ms poststimulus (Fig. S4).

Using this method, single-trial variability (STV) of temporally specific EEG components captured distinct neural substrates following standard and target stimuli at different stages of the reorienting response ^27,38,39^ (see scalp distributions in Fig. S4). We then used these EEG components to inform fMRI analysis, by building a general linear model (GLM) where STVs of EEG components were used to create BOLD predictors (Fig. S5). Specifically, EEG STV regressors’ onset were aligned to different poststimulus windows, with the regressors’ amplitude parametrically modulated by standard and target trials’ EEG STVs at the corresponding windows. Importantly, in the GLM, traditional regressors (unmodulated stimulus-locked regressors and duration modulated RT regressor) were included to absorb stimulus and behavioral response relevant fluctuations in the BOLD. This design ensured that the clusters we identified are correlated with trial-by-trial variability in EEG components, with the average stimulus-evoked and response-related effects controlled for.

This EEG-informed fMRI analysis revealed a coordinated participation of various cortical regions during different stages of the reorienting response (Fig. 2). From 225 to 375 ms, target EEG STVs covaried with BOLD activity in regions such as the precentral gyrus, postcentral gyrus, and superior parietal lobule. Later into the trial from 375 to 600 ms, target EEG STVs correlated with BOLD signal in regions such as the lateral occipital cortex, medial prefrontal cortex, supplementary motor area, insular cortex, and the operculum cortex.

**Figure 2.**
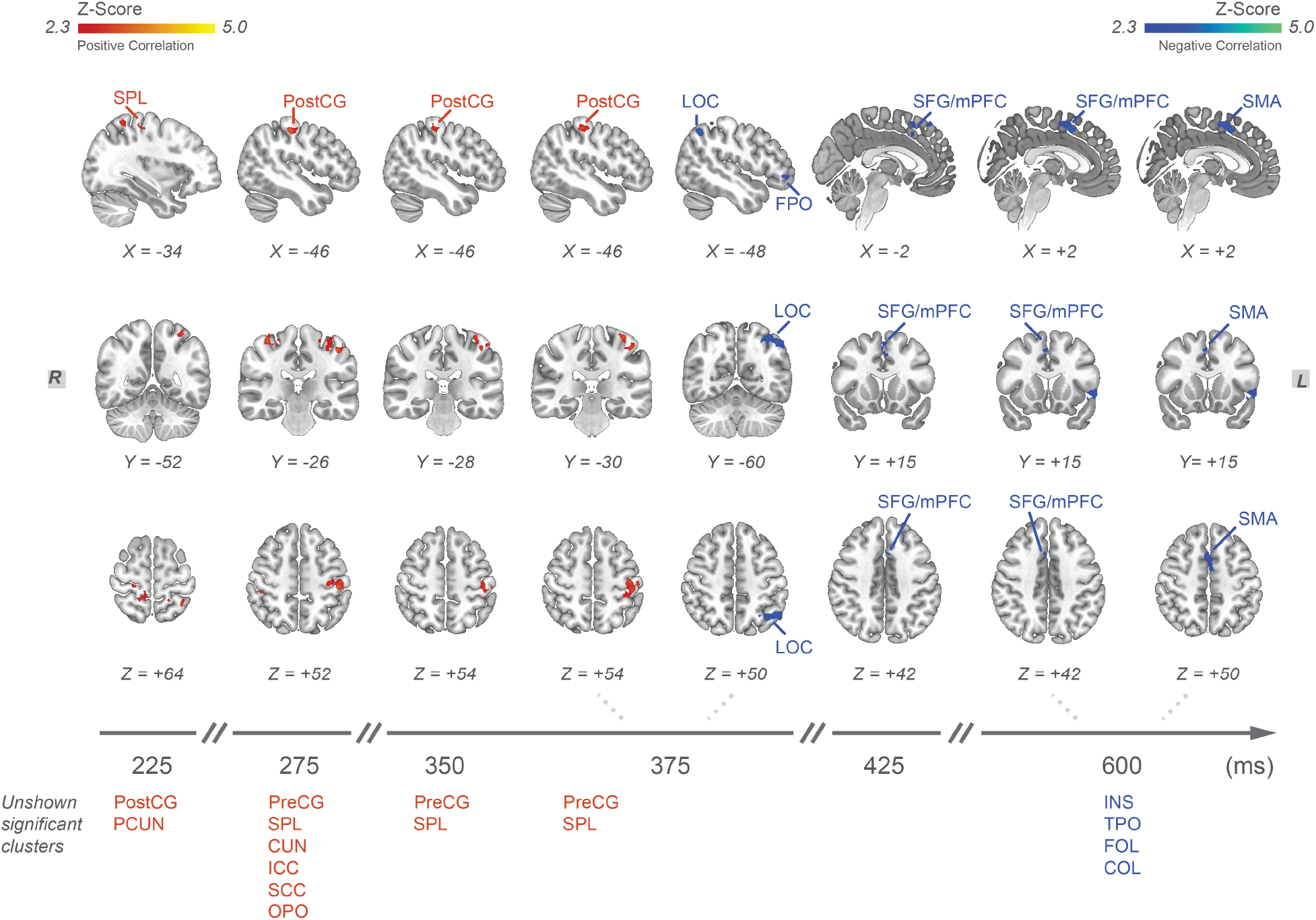
Evolution of activity in cortical regions whose BOLD covary with task-relevant STVs of EEG components. Activation maps showing spatial correlates of target STVs of EEG components at different poststimulus windows. All results were multiple comparisons corrected with *z* > 2.3 and *P* < 0.05. X, Y, Z are MNI coordinates. Positive and negative effects are highlighted in warm and cool color, respectively. Significant clusters not shown in the maps are listed at the bottom. PreCG, precentral gyrus. PostCG, postcentral gyrus. SPL, superior parietal lobule. PCUN, precuneous cortex. CUN, cuneal cortex. ICC, intracalcarine cortex. SCC, supracalcarine cortex. OPO, occipital pole. LOC, lateral occipital pole. FPO, frontal pole. SFG, superior frontal gyrus. mPFC, medial prefrontal cortex. SMA, supplementary motor area. INS, insular cortex. TPO, temporal pole. FOL, frontal operculum cortex. COL, central operculum cortex. PU, putamen. WM, white matter.

**Figure S3. Related to Figure 2.**
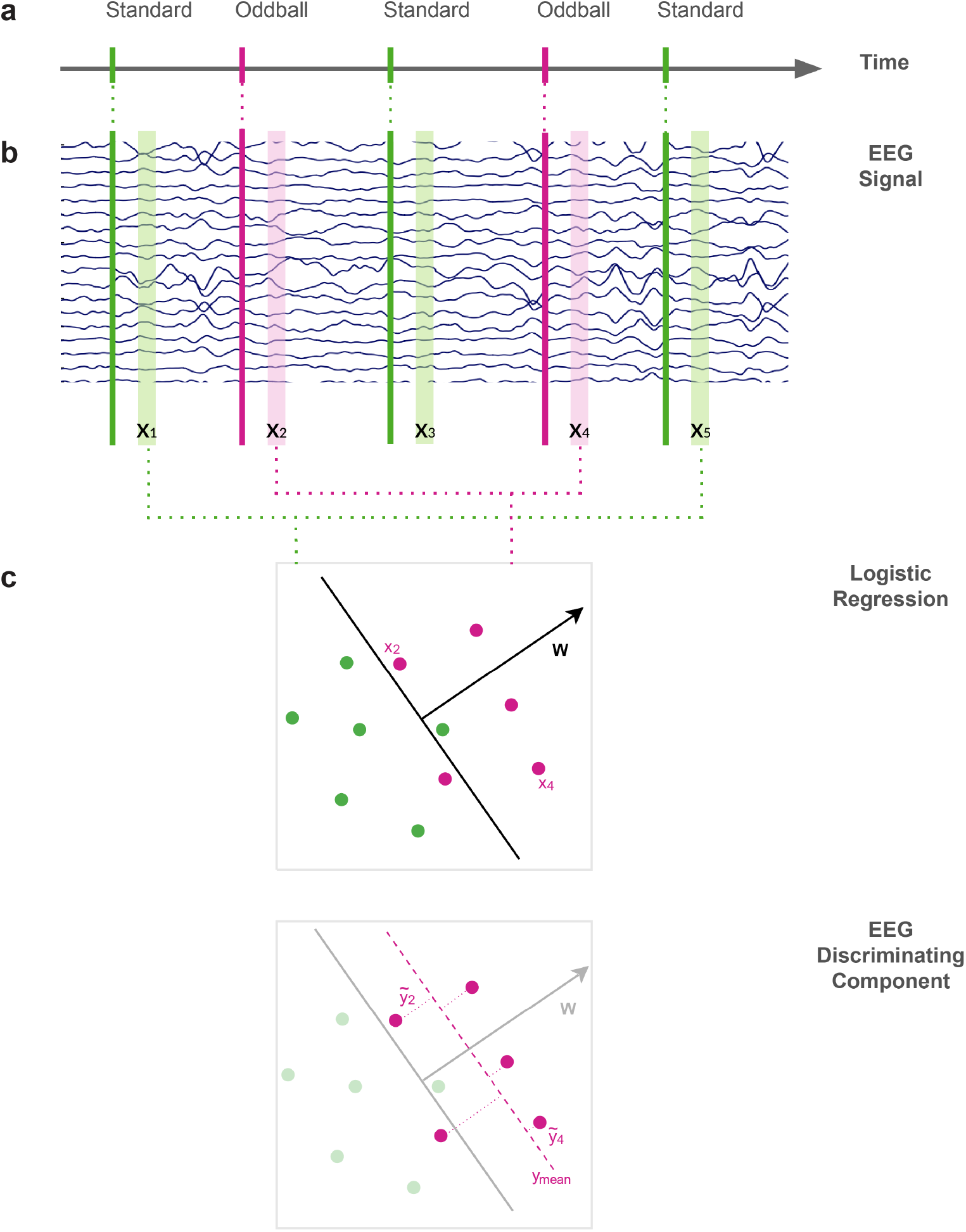
Illustration of single-trial EEG analysis. (**a**) Onsets for a series of standard (green) and oddball (pink) stimuli. (**b**) Specific temporal windows (vertical shaded regions) were used for EEG signal extraction. All time windows had a width of 50 ms and the window center was shifted from 0 ms to 1000 ms relative to stimulus onset, in 25 ms increments. (**c**) Logistic regression was used to create the EEG discriminating component. Specifically, the goal of this algorithm was to find a set of weights *w* that achieves maximal discrimination between multidimensional EEG signal *x*_*i*_ of target versus standard trials within each time window. Applying *w* to *x*_*i*_ results in a projection *y*_*i*_, which captures the distance to the decision boundary for each trial.

**Figure S4. Related to Figure 2.**
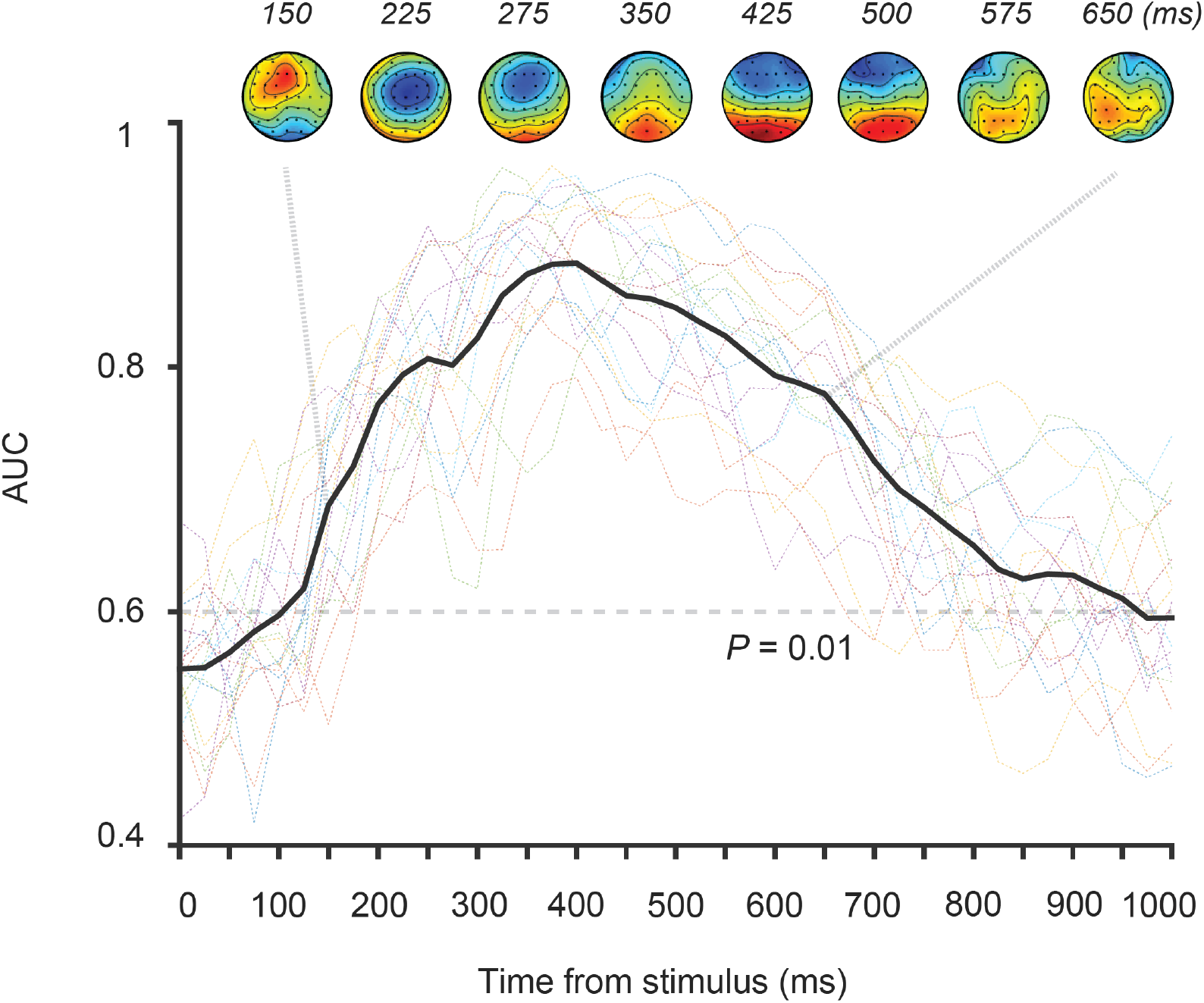
Scalp distributions (i.e., forward models) of temporally specific EEG components were shown from 150 to 650 ms poststimulus (top panel). Group level classifier performance indicated by area under the ROC curve (i.e., AUC) was also shown (bold black line). All panels: group average (*N* = 19); colored dotted lines, individual subjects; dashed line, AUC value with *P* = 0.01; statistics, permutation test.

**Figure S5. Related to Figure 2.**
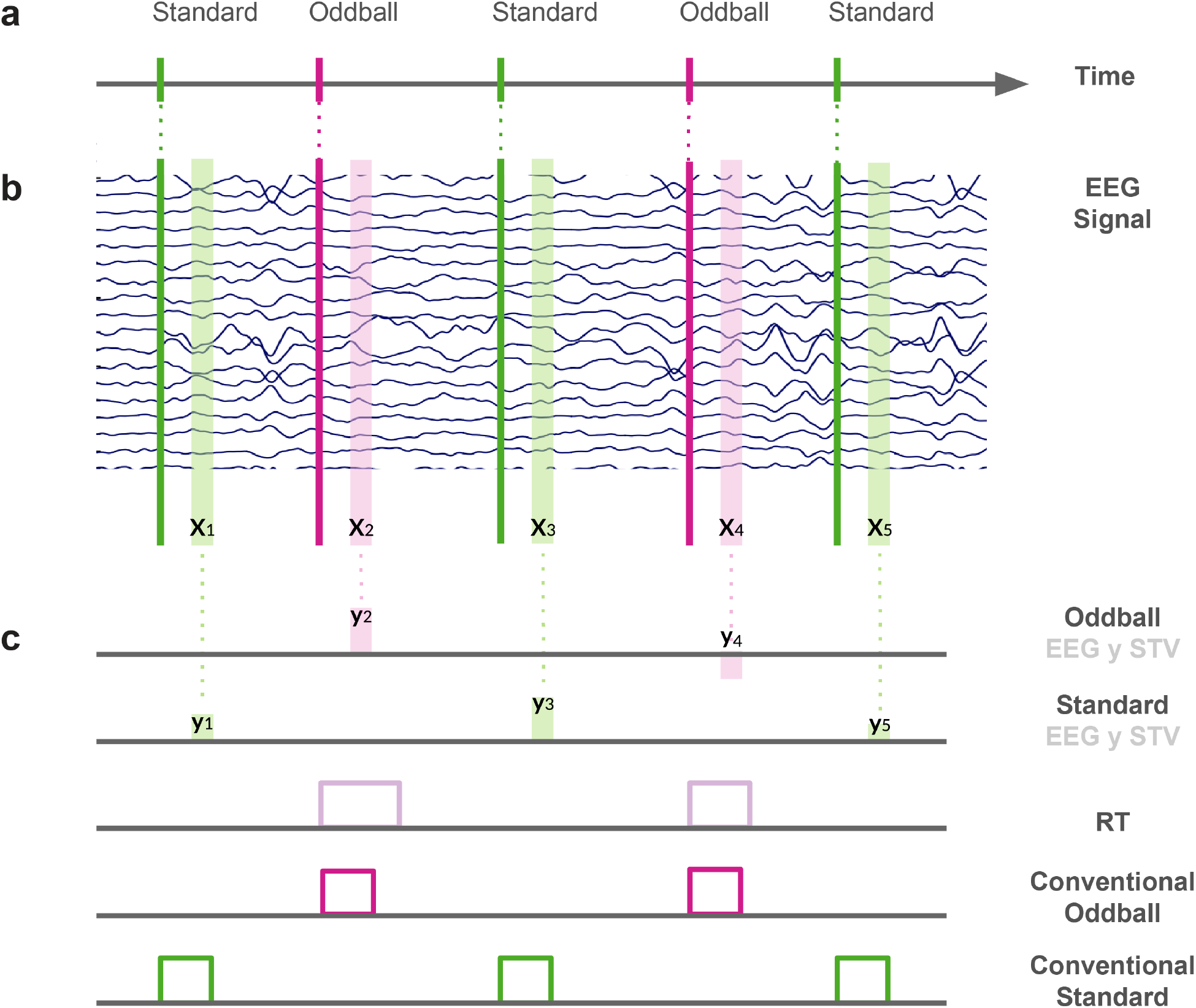
Five regressors were constructed in the EEG-informed fMRI analysis. See section “Whole-brain voxel-wise analysis: EEG-informed fMRI analysis” in Methods for details.

### Fluctuations in evoked and baseline pupil diameter correlated with variability in early and late task-relevant processes

After identifying the temporal cascade of cortical activity related to reorienting using EEG-informed fMRI analysis, we turned to the main goal of this work: investigation of interactions between arousal and reorienting. Specifically, we treated spatial correlates of target EEG components at different time windows as cortical ROIs, and extracted BOLD signal from each ROI. We then used BOLD from each ROI to predict BPD and TPR of target trials with a GLM. This second GLM was constructed to examine the extent to which pupil-linked arousal can be explained by BOLD signal characterizing different task-relevant processes.

We found that when modeling BPD, BOLD signal corresponding to EEG components at 275 and 425 ms contributed significantly. Specifically, BOLD signal corresponding to EEG component at 275 ms negatively contributed to the prediction, whereas BOLD corresponding to EEG component at 425 ms positively contributed to the prediction (GLM group level coefficient estimate at 275 ms window: *b* = -0.106, *P* = 0.002; at 425 ms window: *b* = 0.039, *P* = 0.008, Fig. 3a). Meanwhile, when modeling TPR, BOLD corresponding to EEG components at 225 and 275 ms contributed significantly. Specifically, BOLD corresponding to EEG component at 225 ms negatively predicted TPR, whereas BOLD corresponding to EEG component at 275 ms positively predicted TPR (GLM group level coefficient estimate at 225 ms window: *b* = -0.068, *P* = 0.002; at 275 ms window: *b* = 0.063, *P* = 0.038, Fig. 3b).

**Figure 3.**
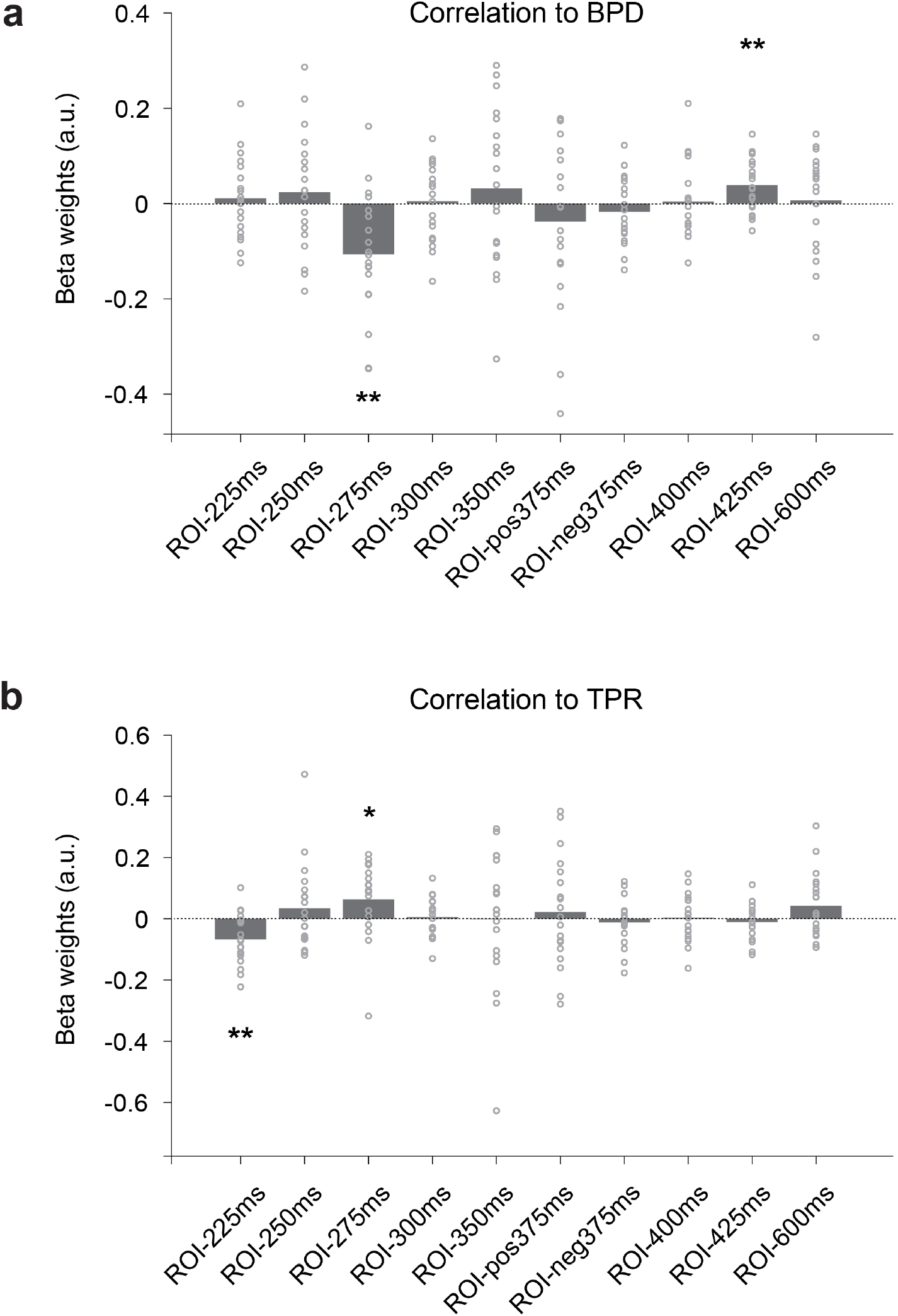
Predict baseline and evoked pupil diameter using ROI-extracted BOLD. (**a**) Regression weights for the linear relationship between ROI-extracted BOLD and BPD. (**b**) is similar to panel (**a**), but for predicting TPR. Time series extracted from ten ROIs were used as regressors in a single GLM. The ROIs were identified from EEG-informed fMRI analysis (e.g., ROI-225ms corresponds to the significant clusters identified by target STVs of EEG component at 225 ms). See section “ROI-based fMRI analysis: modeling pupil with results from EEG-informed fMRI analysis” in Methods for details. All panels: group average (*N* = 19); data points, individual subjects; statistics, one-sample t-test. ** *P* < 0.01, * *P* < 0.05, uncorrected.

### Activity at the core of LC covaried with baseline pupil diameter and cortical activity at P3 latency

Since the level of arousal has been linked to both pupil diameter and LC activity, we additionally investigated if cortical activity associated with pupil-linked arousal are also associated with functional BOLD signal extracted from the core of the LC (for brevity, “the core of the LC” is referred to as “the LC” in subsequent text. See Fig. 4 and section ***ROI-based analysis: LC delineation*** in **Methods** for descriptions and illustrations of LC delineation). Specifically, analogous with EEG-informed fMRI analysis where we used stimulus and EEG STV regressors to predict voxel-based BOLD in the cortex - here we used the same explanatory variables to predict BOLD in the LC. We found that at the three time windows (225, 275, and 425 ms) where cortical activity exhibited significant correlations with pupillary measures, target EEG STVs at neither of the three windows were significantly correlated with BOLD at LC (GLM group level coefficient estimate for EEG STV of target trials at 225 ms window: *b* = 0.015, *P* = 0.108; at 275 ms window: *b* = 0.016, *P* = 0.121; at 425 ms window: *b* = 0.011, *P* = 0.196, Fig. 5a-c). On the contrary, at windows which were in close temporal proximity (i.e., 250 and 300 ms) to the previously examined windows, target EEG STVs were significantly correlated with BOLD at LC (GLM group level coefficient estimate at 250 ms window: *b* = 0.019, *P* = 0.029; at 300 ms window: *b* = 0.026, *P* = 0.022, Fig. 5d-e).

**Figure 4.**
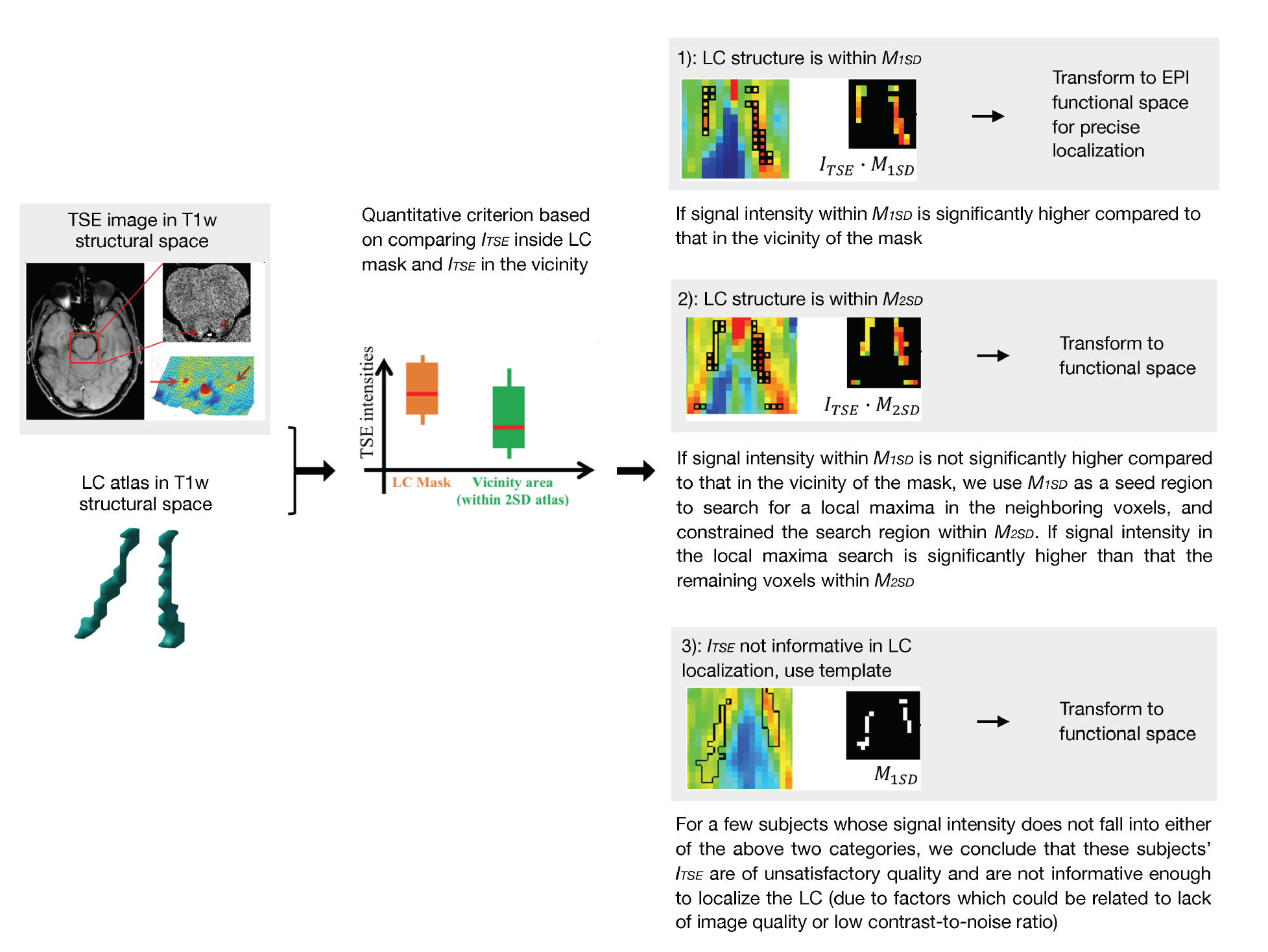
LC delineation. Using predefined LC atlas and acquired TSE images to delineate LC. See section “ROI-based analysis: LC delineation” in Methods for details. I_TSE_, intensity of TSE image. M_1SD_, one standard deviation LC mask. M_2SD_, two standard deviations LC mask.

**Figure 5.**
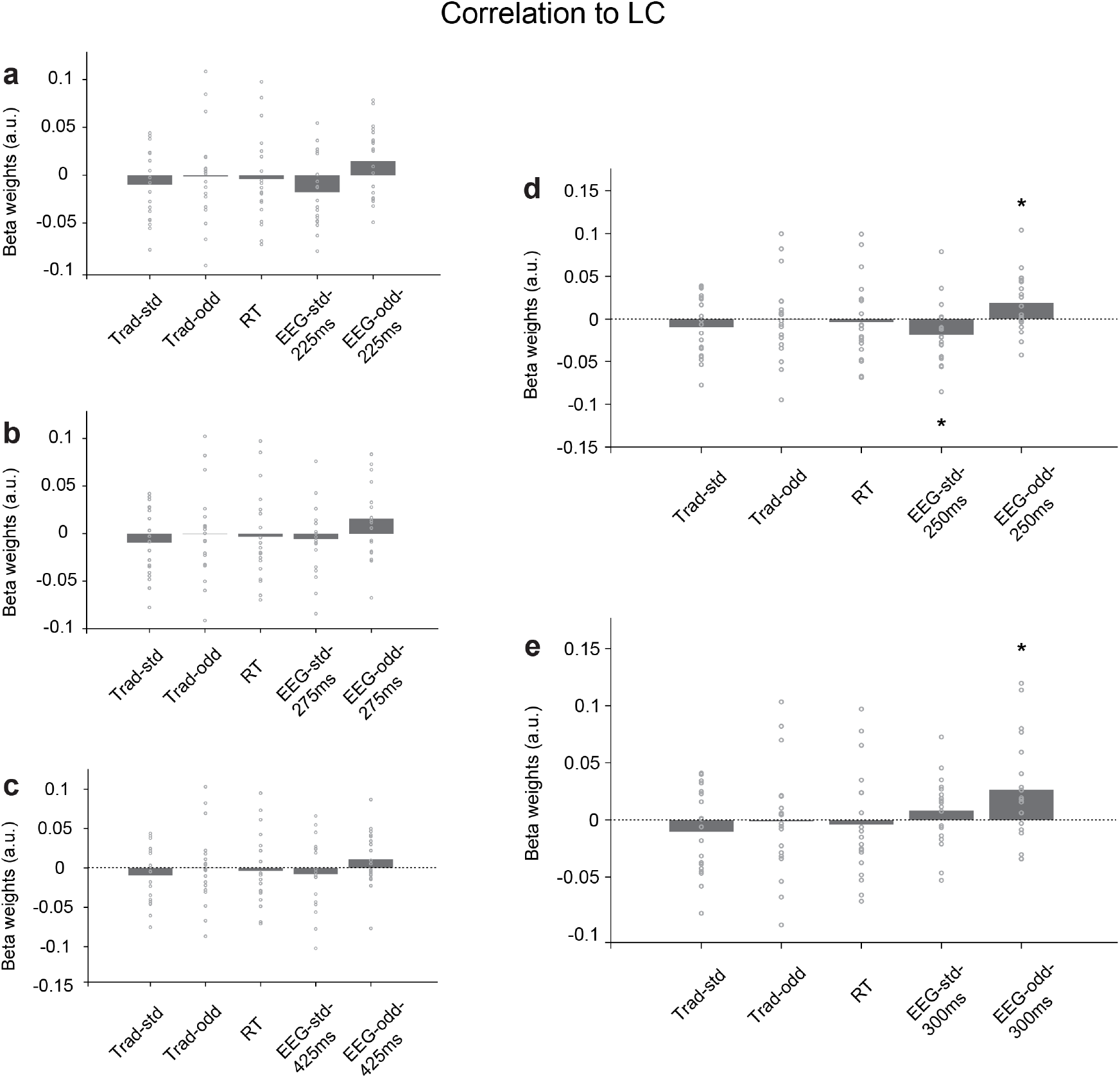
Predict BOLD at LC with STVs of EEG components. (**a**) Regression weights for the linear relationship between BOLD at LC, and traditional and EEG STV regressors at 225 ms. (**b-e**) are similar to panel (**a**), but used EEG STV regressors at 250 ms (**d**), 275 ms (**b**), 300 ms (**e**) and 425 ms (**c**). A total of five regressors were used in each regression: two regressors modeling the effect of standard and oddball (target) stimulus (Trad-std and Trad-odd, respectively); one regressor modeling the effect of behavioral response (RT); and two regressors modeling the effect of temporally specific standard and oddball EEG components (EEG-std- and EEG-odd-at certain temporal window, respectively). All panels: group average (*N* = 19); data points, individual subjects; statistics, one-sample t-test. ** *P* < 0.01, * *P* < 0.05, uncorrected.

Lastly, in order to investigate the extent to which pupil diameter relates to LC activity, we used both pupillary measures (BPD and TPR) to model functional activity at the LC. We found that independent of the stimulus type, there was a robust relationship between trial-by-trial variability of BPD and fluctuations of BOLD at LC (GLM group level coefficient estimate for BPD: *b* = -0.034, *P* = 0.005, Fig. 6a). This relationship was mainly driven by the standard trials, with the target trials exhibiting a similar trend (GLM group level coefficient estimate for BPD of standard trials: *b* = -0.026, *P* = 0.008; for BPD of target trials: *b* = -0.013, *P* = 0.453, Fig. 6b).

**Figure 6.**
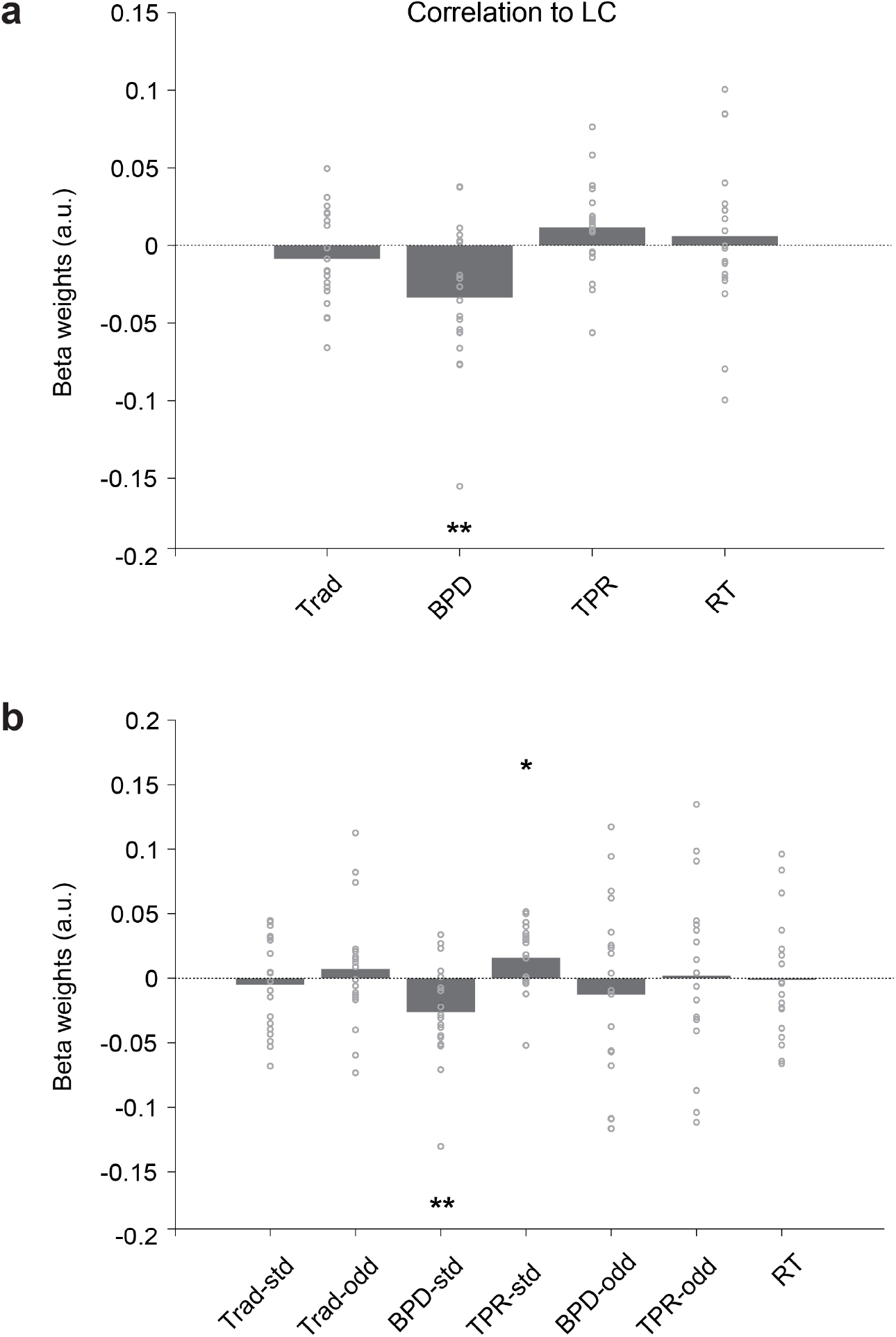
Predict BOLD at LC with baseline and evoked pupillary measures. (**a**) Regression weights for the linear relationship between BOLD at LC and stimulus type-independent pupillary measures. (**b**) is similar to panel (**a**), but using stimulus type-dependent pupillary measures. A total of four regressors were used in (**a**): one regressor modeling the effect of stimulus (Trad); two regressors modeling the effect of pupil (BPD and TPR); and one regressor modeling the effect of behavioral response (RT). In (**b**), the stimulus and pupil regressors were split per stimulus type (-std and -odd), yielding a total of seven regressors. All panels: group average (*N* = 19); data points, individual subjects; statistics, one-sample t-test. ** *P* < 0.01, * *P* < 0.05, uncorrected.

## Discussion

The goal of this study was to investigate the spatiotemporal dynamics between pupil/ LC-linked arousal and attention reorienting. Specifically, we integrated pupillometry, EEG, and fMRI to investigate how different levels of tonic and phasic arousal relate to cortical activity at various stages of the reorienting response. This was achieved by first separating cortical activity associated with reorienting across time and space with the EEG-informed fMRI analysis, and then correlating these spatiotemporally specific activity to pupillary measures and LC responses to relate cortical dynamics to pupil/LC-linked arousal. We found that while tonic and phasic pupil-linked arousal (indexed by BPD and TPR, respectively) were distinctly associated with certain cortical activity at different poststimulus times, both modes of arousal were linked to the same process at 275 ms poststimulus. Additionally, LC-linked arousal (indexed by functional BOLD responses at LC) was also related to cortical activity at multiple poststimulus temporal windows (at 250 and 300 ms) which were in the temporal vicinity of the windows where significant correlations were observed between pupillary measures and task-relevant cortical activity.

### Inverse relationship between phasic pupil-linked arousal and 225 ms cortical activity is suggestive of reduction of response inhibition for target reorienting trials

For the EEG discriminating component at 225 ms, we found its BOLD STVs to be negatively correlated with TPR (Fig. 3b). Given the timing, scalp distribution (frontocentral negativity as illustrated in Fig. S4), and BOLD spatial correlates (PostCG and SPL as illustrated in Fig. 2), this component likely captures neural processes related to the N2 ERP component. The N2 is a negative-going wave which typically peaks between 200 and 350 ms poststimulus, and is most prominent over the frontocentral sites when elicited by auditory stimuli ^40,41^. This second major negative peak after the stimulus onset includes a family of different responses such as the N2a (or mismatch negativity (MMN)) and N2b (or anterior N2), and is therefore linked to a number of neural processes such as the detection of mismatch and cognitive control ^40-42^. Specifically, the relatively automatic MMN is elicited when an auditory stimulus differs from its preceding stimuli, even if the stimulus is task-irrelevant or not attended. The anterior N2, on the other hand, can be evoked by the same auditory mismatch, but only when the stimulus is task-relevant or attended; this effect is also sensitive to response inhibition, and is especially pronounced when motor response is withheld ^40-42^. In the context of an auditory oddball paradigm, subjects are instructed to ignore the standard sound while attending and responding to the target sound. The EEG discriminating component at 225 ms therefore captures both the relatively preattentive MMN and the voluntary attention driven anterior N2 evoked by an infrequent task-relevant target stimulus. Spatial correlates of this EEG component at PostCG and SPL further suggests the involvement of the frontoparietal attention network and primary sensory and motor areas.

While MMN cannot be easily isolated from anterior N2 due to the significant amount of overlap between these two subcomponents ^40,41^, both responses are associated with mismatch detection. These activity are likely to increase in light of heightened phasic pupil-linked arousal, resulting in a positive correlation between TPR and BOLD STVs of this 225 ms EEG component. What inferences can we make then, on the negative link between TPR and BOLD spatial correlates of this component?

One explanation may be that under task-relevant target conditions, increased phasic pupil-linked arousal has an overpowering effect in lifting response inhibition versus detecting mismatch (as the 225 ms EEG component is involved with both). Since BOLD STVs only come from target trials where subjects were instructed to make instead of withhold motor responses, the fact that increased TPR is associated with decreased BOLD in target trials can be interpreted as heightened phasic pupil-linked arousal is related to weaker response inhibition (or stronger promotion of task-relevant behavior). While the positive relationship between TPR and mismatch detection (associated with MMN and portions of anterior N2) and the negative relationship between TPR and response inhibition (associated with other portions of anterior N2) cannot be separately estimated, the overall negative correlation between TPR and BOLD STVs of the 225 ms component indicates a stronger contribution from the response inhibition relevant neural processes.

This observation allows us to make further speculations on how heightened phasic pupil-linked arousal may facilitate the brain’s transition from a long-standing state to a transient state in light of task-relevant information (see another effective connectivity work from our group which provides an in-depth investigation on the role of phasic pupil-linked arousal in network reset ^43^). With infrequent target stimuli sporadically interspersed among frequent standard stimuli, the brain is tuned to withhold motor responses under most conditions. However, under the combined influence of increased phasic pupil-linked arousal, automatic as well as voluntary attention directed to the incoming salient target stimulus, response inhibition is likely lifted to facilitate the execution of motor response which is relevant to the task. Three sets of previous findings support this claim: (i) the established understanding that the LC-NE system affects neuronal activity in the somatosensory cortex by suppressing spontaneous activity more than transient sensory-evoked responses (see review ^44^); (ii) the demonstrated causal relationship between microstimulation of LC and pupil dilation ^13^; and (iii) the reported connections between phasic arousal and the execution not withhold of motor responses in animal work ^45,46^. While pupil dilations have been linked to the behavioral response in previous studies, investigations have mostly been confined to the relationship between amplitudes of pupil dilation and RT (which serves as a generic index of task performance) ^18,24,25,47^. In this study, by capitalizing on the modality specific spatiotemporal information extracted from simultaneous pupillometry and EEG-fMRI recordings, we were able to uncover the relationship between TPR and another measure which is related with the response: a spatiotemporally specific component representative of the inhibition of response. Our findings support previous studies which have linked TPR to both bottom-up and top-down processes (see review ^21^). To the best of our knowledge, the present study also provides the first demonstration in humans of the inverse relationship between phasic pupil-linked arousal and reduction of response inhibition.

### Phasic and tonic pupil-linked arousal engage cortical networks at the beginning of the classic P3 response

The task-relevant component whose BOLD STVs were associated with both tonic and phasic pupillary measures was temporally situated at 275 ms poststimulus (Fig. 3). This time sits at what is usually considered the beginning of the P3 (250 to 500ms) elicited in an oddball paradigm ^18,24,25,48^. Additionally, spatial correlates of this 275 ms EEG component were identified at PreCG, PostCG, SPL and various regions in the occipital lobe (ICC, SCC, and OPO). Taken together, activation of the frontal, parietal and occipital lobes at a P3 time suggest involvement of the frontoparietal attention network and primary sensory areas in stimulus-driven attention reorienting and goal-driven task performing ^1,48^.

Additional inferences can thus be drawn from the observed relationship between pupillary measures and BOLD STVs of this component. Specifically, our analysis revealed the presence of a positive correlation between TPR and BOLD, and a negative correlation between BPD and BOLD. This finding suggests that increased TPR and decreased BPD were associated with increased BOLD of this component. The positive correlation between TPR and BOLD provides additional evidence supporting the proposed link between phasic pupil-linked arousal and task-relevant cognitive processes in the temporal vicinity of P3 ^1,8,48^. The negative correlation between BPD and BOLD, on the other hand, supports the proposed negative effect of tonic pupil-linked arousal on task-relevant responses (that is, reduced performance in light of increased tonic LC activity) ^8,49^. Specifically, BPD has been robustly linked to P3, with elevated levels of BPD associated with diminished amplitudes of P3, and vice versa ^18,24,25^. Along with previous findings, our results suggest that a high level of tonic pupil-linked arousal decreases the amplitude of task-relevant responses such as the P3. This supports the hypothesis that task-relevant performance would be optimal under intermediate level of tonic arousal ^8,49^.

### Tonic pupil-linked arousal correlates with cortical networks engaged at 425 ms which is suggestive of a switching from exploitation to exploration

For the task-relevant component at 425 ms, its BOLD STVs were positively associated with BPD, the tonic pupillary measure (Fig. 3a). This finding suggests that increased BPD is associated with increased BOLD of this component which is spatially localized at SFG and mPFC. Together with our observations of a significant negative correlation between stimulus type-independent BPD and the activity at LC (Fig. 6a), these findings suggest that BPD could be tied to the behavior of LC. Specifically, when returns of task performance no longer surpasses the investments to the performance based on frontal structures’ evaluation, LC switches to a tonic mode where baseline activity is elevated and phasic activity becomes absent. The tonic activity of the LC-NE system thus favors exploration of other potentially rewarding behaviors over exploitation of performance of the current task ^8^. It is therefore likely that at least to some extent, our results reflect the positive impact of tonic pupil-linked arousal on exploration over exploitation.

### Pupil diameter and task-evoked LC responses capture both common and unique cortical activity

Modeling pupil with results from EEG-informed fMRI analysis allowed us to evaluate and infer the relationship between pupil-linked arousal and temporally specific cortical activity related to reorienting. Meanwhile, quantifying task-evoked LC responses with EEG components enabled further investigation on how LC-linked arousal relates to neural processes at different temporal stages of the reorienting response. These two types of analyses complement one another in terms of the specific aspect of global arousal that was being captured (i.e., pupil-linked arousal versus changes in cortical arousal modulated by task-evoked LC responses).

For instance, while BPD and TPR were related to BOLD STVs of EEG components at certain poststimulus windows (225, 275, or 425 ms), LC activity did not covary with EEG components at those specific windows (Fig. 5a-c). On the contrary, the activity at LC correlated with EEG components in close temporal proximity (i.e., 250 and 300 ms, Fig. 5d-e) to the aforementioned windows which exhibited significant correlations with pupillary measures. These observations suggest that while there is common neural activity measurable by pupil diameter and functional BOLD responses at LC, each modality-specific measure also captures unique information. In support of this claim, increasing evidence suggests that the relationship between LC activity and pupil diameter is more heterogeneous than homogenous (see reviews ^21,44^, and a recent study capturing the dissociation between pupil diameter and LC spiking activity ^50^). In our study, such distinction is best characterized by the relationship between pupil- or LC-linked arousal and EEG components at 225 and 275 ms. Specifically, BPD and TPR exhibited an opposite relationship with BOLD spatial correlates of the EEG components at 225 and 275 ms (Fig. 3), which in turn showed no significant correlation with the LC activity (Fig. 5a-b). On the contrary, at 250 and 300 ms where the relationship between pupillary measures and BOLD STVs of the EEG components were comparable and of the same sign, the EEG components showed significant correlation with the LC activity (Fig. 5d-e). These observations illustrate the unique contribution of including both pupil and LC measures in investigating how arousal relates to neural processes involved in attention reorienting. If not investigated separately, the opposite yet distinct relationship between the two pupillary measures and neural processes at 225 and 275 ms may be canceled out. Such inferences would not be possible in cases where modalities such as pupillometry, EEG, or fMRI were acquired separately, and further demonstrates the advantage one could gain from simultaneous multi-modal acquisitions.

### Limitations and future directions

In our study, we combined pupil, EEG, and fMRI data in the time domain with a hierarchical, and asymmetric approach. Specifically, we (i) extracted modality-specific measures from the time domain; (ii) examined the covariability underlying EEG and fMRI with an asymmetric fusion method (i.e., EEG-informed fMRI analysis); and (iii) used results from the EEG-fMRI analysis to model pupillary measures and LC responses. Our results therefore captured the correlational, but not causal relationship between cortical dynamics and pupil/LC-linked arousal. Future work using effective connectivity can aid in the identification of directionality in these observed interplays.

In addition, while our analysis focused on the time domain and used an asymmetric approach, symmetric or frequency-domain based fusion methods could also contribute to a more comprehensive understanding of the relationship between latent neural states and arousal. A notable example of symmetrical pupil-EEG-fMRI fusion was presented in Groot et al., 2021 ^51^. The authors extracted features from all three modalities and used a support vector machine to combine features, and to investigate the neural signature of task unrelated thoughts. Future investigations are likely needed to reveal the strengths of each fusion method in revealing neural dynamics in various tasks when pupil, EEG, and fMRI are acquired simultaneously.

Another possibility of analyzing this dataset resides in the frequency domain. In the only two reported simultaneous pupillometry and EEG-fMRI studies, Groot et al., 2021 ^51^ and Mayeli et al., 2020 ^52^ tested if frequency-domain features can be reliably tied to vigilance or task unrelated thoughts, respectively. Mayeli et al., 2020 ^52^ argued that frontal and occipital beta power can serve as a reliable index for vigilance level of healthy humans during a resting state MR scan. Considering the much investigated relationship between spectral oscillations and an abundance of cognitive processes (e.g., cortical idling ^53^, mind wandering ^54^, inhibition of task-irrelevant regions ^55^, and information processing ^56^), future simultaneous pupillometry and EEG-fMRI studies are needed to decipher the rich inferences embedded in relationships between spectral oscillations and arousal relevant processes.

### Unique contributions of a simultaneous pupillometry and EEG-fMRI study

Compared with acquiring EEG or fMRI separately, simultaneous acquisition of EEG and fMRI has less popularity. This is in part due to the technical challenges involved in concurrent data acquisition, and partly due to the scarcity of effective data fusion methods ^39^. Understandably, relative to simultaneous EEG-fMRI, due to the increased technical and methodological hurdles involved with the addition of a third modality, concurrent acquisitions combining three modalities (pupillometry, EEG, and fMRI) have so far been applied in very few instances. In fact, to the best of our knowledge, simultaneous acquisition of pupillometry and EEG-fMRI has not been reported until the recent two years, and so far has only been discussed in two studies ^51,52^. In this paper, we contribute to this burgeoning conversation and aim to demonstrate that despite the challenges, concurrently recorded pupillometry and EEG-fMRI data offer unparalleled potential in revealing spatiotemporal dynamics of cortical networks.

Previous works investigating the relationship between the arousal and reorienting systems typically take well-studied neural activity in the reorienting response (such as P3) and examine how different levels of arousal affect such processes. In our study, we did not confine ourselves to established or widely examined responses prior to the analysis. Rather, we first used a data-driven approach to extract temporally specific task-relevant EEG components which had millisecond resolutions. We then used an EEG-informed fMRI analysis to examine spatial correlates of these components which represent specific cortical activity. By correlating pupillary measures with results from the EEG-fMRI analysis, we investigated how certain cortical activity (in a specific time frame and related to specific regions) interacted with different levels of tonic and phasic pupil-linked arousal. With these novel approaches, we found that different modes of pupil-linked arousal interact with cortical activity at different temporal stages of the reorienting response. Specifically, phasic pupil-linked arousal relates to the reduction of response inhibition, while tonic pupil-linked arousal relates to the brain’s preference of exploration over exploitation when task utility wanes.

Notably, our work is of value to studies in both nonclinical and clinical settings. For studies involving nonclinical populations, future works can use the technique described in this study to systematically investigate how arousal interacts with reorienting in a more complex setting (e.g., in a virtual reality environment while harvesting high spatiotemporal resolutions). Meanwhile, this study can facilitate treatment development for clinical conditions related with ineffective attention reorienting (e.g., attention deficit hyperactivity disorder, autism, depression and post-traumatic stress disorder (PTSD)). For instance, if populations of a certain clinical condition tend to favor exploitation over exploration (e.g., prolonged sustained attention to threat in PTSD), their baseline pupil diameter can be monitored to index deterioration or improvement of their conditions.

In summary, this study provides new evidence which reveals unique relationships between tonic and phasic pupil-linked arousal and attention reorienting in the human brain in a target detection task. These findings are enabled by a suite of multi-modal acquisition and analysis approaches which are capable of revealing comprehensive spatiotemporal dynamics in cortical and subcortical networks.

## Acknowledgements

The authors wish to thank Josef Faller, Ray Lee, and members of the Columbia Magnetic Resonance Research Center for their expert assistance in data acquisition. This work was supported by an Army Research Laboratory Cooperative Agreement (grant W911NF-10-2-0022 to P.S.), a Vannevar Bush Faculty Fellowship from the US Department of Defense (grant N00014-20-1-2027 to P.S.), and a seed grant for MR Studies Program of the Zuckerman Mind Brain Behavior Institute at Columbia University (grant CU-ZI-MRS-0006 to L.H. and P.S.).

## Author Contributions

L.H. and P.S. conceived the study. L.H. developed experimental protocol and collected data. L.H. and H.H. analyzed the data. L.H., H.H. and P.S. discussed the analyses. L.H. wrote the initial draft of the manuscript. H.H. and P.S. reviewed and edited the manuscript. L.H. and P.S. acquired funding.

## Methods

### Experimental Model and Subject Details

Twenty-five human subjects participated in this study. Six were excluded from further analysis due to incomplete data (*N* = 2), abnormality in collected data (*N* = 2), excessive movement (*N* = 1) and inability to complete the task per instruction (*N* = 1), respectively. The remaining 19 subjects (6 males) aged between 18 to 32 years (mean age 25.9 years) were included in all subsequent analyses. All participants had normal or corrected-to-normal vision and no reported history of psychiatric or neurological disorders. The Columbia University Institutional Review Board approved this study, and written informed consent was obtained from every participant prior to the experiment.

### Method Details

#### Behavioral paradigm

We used an auditory oddball paradigm with 80% of standard and 20% of target (oddball) stimuli in this study (Fig. 1a). The standard stimuli were pure tones with a frequency of 350 Hz, while the target stimuli were broadband (“laser gun”) sounds. All stimuli had a duration of 200 ms with an inter-trial interval sampled from a uniform distribution between 2 s and 3 s.

We created and presented the paradigm using PsychToolbox in MATLAB. Auditory stimuli were delivered to subjects via earphones. Throughout the experiment, a fixation target was presented on a gray background for the subjects to fixate. This specific type of fixation target has been shown to outperform other alternatives in lowering dispersion and microsaccade rate ^62^, thus facilitating stable recording of subjects’ pupil diameter. The fixation target was presented to a screen placed outside the scanner bore, and viewed by the subjects through a mirror mounted on the head coil.

Before the main experiment in the scanner, subjects were first familiarized with the paradigm through a short practice run of the task outside of the scanner. Once inside the scanner, they were instructed to maintain fixation to the fixation target, and to respond to the target stimuli as quickly and accurately as possible, by pressing a button on a MR-compatible response box with their right index finger. Every subject was scheduled to complete 5 runs of 105 trials each. On average, subjects completed 3 to 5 runs (4.7 ± 0.7 runs, mean ± SD; SD: standard deviation). A randomized trial order was maintained with two constraints: (i) the first five trials of each run were constrained to be standards, so that the subjects were well settled into the experiment before the appearance of the first target stimulus; and (ii) no consecutive target trials were allowed, so that enough time elapsed for subjects’ pupil diameter to go back to baseline before the onset of another target trial (i.e., inter-target interval was always larger than 4 s).

#### EEG acquisition

We acquired simultaneous pupillometry, EEG, and fMRI inside an MR scanner. Specifically, EEG was collected at a sampling rate of 5 kHz with an MR-compatible EEG amplifier system (BrainAmp MR Plus system, Brain Products). The EEG cap included 64 passive Ag/AgCl electrodes, with 63 electrodes positioned on the scalp according to the international 10-20 system, and 1 electrode positioned on the subject’s back for electrocardiogram (ECG) monitoring. To ensure subject safety during simultaneous EEG and fMRI acquisition, the scalp and ECG electrodes were embedded with series resistors of 10 kOhm and 20 kOhm, respectively. During the experiment, electrodes’ impedances were kept under 25 kOhm (including the built-in resistors on each electrode) to minimize noise in EEG acquisition.

#### Pupillometry acquisition

Concurrently with the EEG and fMRI acquisitions, pupillometry was collected at a sampling rate of 1 kHz with an MR-compatible eye tracker (Eyelink 1000 Plus in Long Range Mount, SR Research). The eye tracker was placed outside the scanner bore, close to the rear end of the scanner table where the subject’s head and the EEG amplifiers were situated. At the start of each scanning session, calibration of the eye tracker was performed to ensure accurate tracking of the subject’s pupil.

#### MRI acquisition

MR images were collected on a 3T Siemens Prisma scanner with a 64-channel head/ neck coil. For each subject, functional images were acquired interleaved in 42 axial slices with a T2*-weighted echo-planar imaging (EPI) sequence: echo time (TE) = 25 ms, repetition time (TR) = 2100 ms, flip angle (FA) = 77 degrees, voxel size = 3 mm x 3 mm x 3 mm, field of view (FOV) = 192 mm x 192 mm. A structural image was acquired with a T1-weighted sequence for anatomical co-registration (TE = 3.95 ms, TR = 2300 ms, FA = 9 degrees, voxel size = 1 mm x 1 mm x 1 mm, FOV = 176 mm x 248 mm). To facilitate co-registration between structural and functional images, an additional single-volume EPI image was acquired with a T2*-weighted sequence using a resolution higher than that in the functional EPI images (TE = 30 ms, TR = 6000 ms, FA = 90 degrees, voxel size = 2 mm x 2 mm x 3 mm, FOV = 192 mm x 192 mm). Lastly, to localize LC, neuromelanin-sensitive images were acquired in 10 axial slices with a T1-turbo spin echo (T1-TSE) sequence (TE = 14 ms, TR = 600 ms, FA = 90 degrees, voxel size = 0.4 mm x 0.4 mm x 3 mm, FOV = 166 mm x 205 mm). Specifically, TSE images were acquired in an orientation perpendicular to the plane of the brainstem to facilitate accurate delineation of LC ^12,63,64^.

### Quantification and Statistical Analysis

#### EEG pre-processing

EEG acquired inside the scanner are susceptible to MR environment related artifacts such as the gradient artifacts and ballistocardiogram (BCG) artifacts, due to electromagnetic interactions between the two recording modalities. The pre-processing of EEG data therefore was composed of gradient, BCG, and standard artifacts removal from the EEG signal. To begin with, high amplitude gradient artifacts were removed using an average artifact template subtraction method ^65^. This was performed in a data processing software (BrainVision Analyzer 2, Brain Products) provided by the EEG system manufacturer. Specifically, for EEG acquired simultaneously with each functional MR volume, gradient artifacts from 20 volumes centered on the volume of interest were used to construct an average artifact template, which was subsequently subtracted from the EEG signal. The data was then down-sampled to 500 Hz and filtered by a 10th-order median filter to remove residual spike artifacts. In preparation for BCG artifacts removal, slow drift and high frequency noise irrelevant to neural processes were filtered out by applying a 4th-order bandpass Butterworth filter from 0.5 Hz to 50 Hz on the EEG signal. Filtered EEG data were then concatenated over runs for each subject, and removed of BCG artifacts through the simple mean approach offered in the FMRIB plug-in for EEGLAB ^66,67^. Lastly, the BCG-removed data were re-referenced to common average, and removed of blink artifacts using the independent component analysis methods introduced in EEGLAB ^68^.

Trials were excluded from subsequent analyses based on the following criteria: (i) when probability of data value from a single channel was more than 6 standard deviations from the channel’s mean; (ii) when probability of data value from all channels was more than 2 standard deviations from all channels’ mean; and (iii) when subjects failed to respond to the target stimulus, or incorrectly responded to the standard stimulus. On average, 5% of trials were excluded from each subject.

#### Single-trial EEG analysis

We used a linear discriminant analysis to discriminate single-trial EEG measures between target and standard trials (see Parra et al., 2005 ^35^ and Sajda et al., 2009 ^69^ for an overview of this approach; see Hong et al., 2014 ^19^, Fouragnan et al., 2015 ^37^, Gherman et al., 2018 ^70^, and Franzen et al., 2020 ^38^ for applications of this approach). At various temporal windows within the time span of the trial, we used logistic regression to find a set of spatial weightings that when applied to the multidimensional EEG data, yielded a one-dimensional projection that achieves maximal discrimination between the target and standard conditions. Compared to event-related potential analysis which averages individual-channel measures over trials, this single-trial method not only ensured signal-to-noise ratio (SNR) through spatial integration of multi-channel EEG data, but also preserved trial-by-trial variability which captures task-relevant fluctuation of latent brain states. The resulting one-dimensional projection - also called the EEG discriminating components - were therefore used to identify neural correlates of reorienting.

Specifically, we identified task-evoked discriminating components, by estimating a spatial weighting vector which maximally discriminates EEG data of two conditions at specific temporal windows:

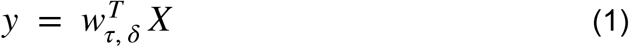

where *τ* is the window center, *δ* is the duration of the window, and 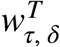 is the transpose of *w*. In practice, this was repeated for 50-ms time windows from 0 ms to 1000 ms relative to stimulus onset, in 25 ms increments. Scalp distributions capturing the mapping from discriminating components to original EEG data were also computed with:

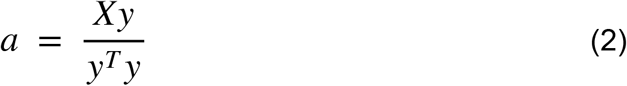

These scalp distributions represent the forward model which can best explain the observed EEG data given the discriminating components which exhibit certain temporal properties (in this case, could maximally discriminate between target versus standard conditions in a specific temporal window).

We quantified the performance of the discriminator at each temporal window by using a leave-one-out approach ^36^ and computing the area under the receiver operating characteristic curve (referred to as Az in subsequent text). To compute a significance level for Az, we used a bootstrapping technique where we randomized the trial labels before performing the leave-one-out test. We repeated this randomization procedure for 100 times to produce a probability distribution for Az, and identified the Az value corresponding to a significance level of *P* = 0.01. Discriminating components at windows whose Az values were significant were used in subsequent analyses.

#### Pupil pre-processing

We adapted the pre-processing pipeline introduced in Urai et al., 2017 ^20^ to prepare pupillometry data for further analysis. Specifically, blinks detected by the manufacturer’s standard algorithms were padded by 150 ms and linearly interpolated. Additional outliers in the remaining time series were detected and removed through a peak detection algorithm. Resulting pupil area data were then converted to pupil diameter and band-pass filtered using a 2nd-order Butterworth filter from 0.01 Hz to 10 Hz to remove slow drift and high frequency noise. Lastly, filtered pupil diameter data for each run was z-scored and down-sampled to 500 Hz to facilitate subsequent multi-modal analyses (i.e., to the same sampling rate of pre-processed EEG data).

#### Pupil analysis

For each trial, prestimulus baseline pupil diameter was extracted as the mean pupil diameter from -500 ms to 0 ms relative to the stimulus onset. Task-evoked pupil response was measured as the maximum deviation from baseline pupil diameter during the two seconds poststimulus.

#### fMRI pre-processing

Pre-processing of the MRI data relied on a number of tools in FSL ^71^. To begin with, we applied bias field correction to functional images to improve the SNR of the BOLD signal. This step was necessary for fMRI data acquired simultaneously with EEG, as the presence of the EEG acquisition system introduced inhomogeneity to the magnetic field and thus could affect the SNR of collected fMRI signal. The specific pre-processing steps therefore included (i) bias field correction by adjusting for variations in spatial intensity using FAST ^72^; (ii) non-brain tissue removal using BET ^73^; (iii) slice-timing correction using Fourier-space time-series phase-shifting; (iv) small head movement correction using MCFLIRT ^74^; and (iv) low-frequency noise removal using a high-pass temporal filtering (Gaussian-weighted least-squares straight line fitting, with sigma = 50 s).

With the pre-processed fMRI data, we performed the subsequent analyses following two separate pipelines: (i) based on individual voxels, by predicting voxel-wise BOLD signal using a set of regressors; or (ii) based on specific regions of interest, by relating BOLD responses from ROIs to physiological or neural measures. The procedures for the delineation of ROIs are described in the sections ***ROI-based fMRI analysis*** below. For voxel-based analyses, functional images were spatially normalized to the standard (MNI) space. This spatial normalization procedure involved using boundary based registration ^75^ to register functional images first to the one-volume high resolution structural image, which was then aligned to the standard space with a combination of linear (FLIRT) and nonlinear (FNIRT) registration approaches ^76-79^. We additionally applied spatial smoothing to functional images in the standard space using a Gaussian kernel of 5 mm full width at half maximum. All other analyses were performed without spatial smoothing.

#### Whole-brain voxel-wise fMRI analysis: overview

We used two approaches to identify brain regions which were either sensitive to the stimulus, or to temporally specific neural processes. These whole-brain voxel-wise statistical analyses were conducted using a general linear modeling (GLM) approach, as implemented in the FEAT tool of FSL. In the GLM, the response variable was the measured BOLD response time series at a given voxel, and the explanatory variable was the design matrix which captured the hypothetical task-evoked neuronal responses. The design matrix was constructed by convolving a set of task-evoked regressors with a canonical hemodynamic response function. Specifically, (i) double-gamma function was used to convolve regressors; (ii) motion parameters generated in the motion correction pre-processing step were included as nuisance regressors; (iii) temporal derivatives of all regressors were included as regressors of no interest; and iv) local autocorrelation correction was performed using the FILM tool ^80^.

The GLM analysis was performed at the subject-level with a fixed effects model, and then at the group-level with a mixed effects model (using FLAME stage 1 ^81-83^). Parameter for each regressor was estimated by minimizing the residual error (i.e., the difference between observed and predicted BOLD responses). The analysis was performed for voxels across the whole brain, and Gaussian random field theory was used to correct for multiple comparisons and to adjust the significance level of identified clusters.

#### Whole-brain voxel-wise analysis: conventional fMRI analysis

With the first approach, we identified brain regions which exhibited distinct average responses after the task-relevant stimulus (i.e., regions with a stronger fMRI response following the target versus the standard stimulus). Specifically, we quantified task-evoked fMRI responses using a set of regressors locked at the time of stimulus. We constructed three boxcar regressors of interest: (i) two unmodulated regressors to model the effect of standard and target stimulus (duration = 0.1 s, height = 1), and (ii) a duration-modulated regressor to control for the effect of behavioral response (onset = target stimulus onset, duration = RT, height = 1, orthogonalized with respect to the target regressor). Significance level of the parameter estimates were determined by thresholding z statistic images using clusters determined by *z* > 3.1, with a corrected cluster significance threshold of *P* = 0.05 ^84^.

#### Whole-brain voxel-wise analysis: EEG-informed fMRI analysis

While the first approach allowed us to identify brain regions that exhibited stimulus-specific responses, the second approach enabled us to assign temporal order to the network of regions that were recruited during reorienting. In particular, we capitalized on the output of our single-trial EEG analysis to build two additional BOLD predictors. At each temporal window where the linear discriminator’s performance was significant, trial-by-trial amplitude of the discriminating component (resulting from equation 1 as illustrated in the ***Single-trial EEG analysis*** section) was used to parametrically modulate the height of boxcar regressors. This GLM therefore included five regressors of interest: (i) three regressors from the conventional fMRI analysis above; (ii) an amplitude-modulated standard EEG STV regressor (onset = window center, duration = 0.1 s, height = of standard trials, orthogonalized with respect to the unmodulated standard regressor); and (iii) an amplitude-modulated target EEG STV regressor (onset = window center, duration = 0.1 s, height = of target trials, orthogonalized with respect to the unmodulated target and duration-modulated RT regressor). Significance level of the parameter estimates were determined by thresholding z statistic images using clusters determined by *z* > 2.3, with a corrected cluster significance threshold of *P* = 0.05 ^84^. As the target EEG STV regressor was the primary regressor of interest, this analysis identified a set of cortical regions whose BOLD covary with task-relevant STVs of EEG components.

#### ROI-based fMRI analysis: overview

While whole-brain analysis enabled us to localize activations bearing specific temporal profiles in attention reorienting, the ROI-based analyses were designed to investigate how pupil-linked arousal or LC activity interacted with these temporally specific cortical activity. In section ***modeling pupil with results from EEG-informed fMRI analysis***, we define significant clusters identified with EEG-informed fMRI analysis as ROIs. By using BOLD from these ROIs to model BPD and TPR, we examined the relationship between pupil-linked arousal and BOLD responses which covary with trial-by-trial variability in temporally specific EEG components. In section ***LC delineation***, we define another ROI which is LC, and describe the techniques we used to delineate this nucleus. In section ***quantifying task-evoked LC responses with EEG***, we use EEG STV modulated regressors to model BOLD responses at LC. This second ROI-based analysis is analogous to the whole brain EEG-informed fMRI analysis, with the former focusing on activity confined to the LC and the latter examining activity across the brain. In addition, in section ***quantifying task-evoked LC responses with pupil***, we describe another ROI-based analysis which evaluated the hypothesis that pupil reflects LC activity. In this analysis, we used both pupillary measures (BPD and TPR) to model functional BOLD signal extracted from LC. All three analyses were conducted using a general linear modeling approach, with parameter estimates computed separately for each subject and tested against 0 across the group.

#### ROI-based fMRI analysis: modeling pupil with results from EEG-informed fMRI analysis

In the first analysis, we modeled task-relevant target pupillary measures with BOLD responses from ROIs identified with EEG-informed fMRI analysis. Specifically, single-trial target pupillary measures (BPD and TPR) were first convolved with a canonical double-gamma hemodynamic response function to prepare the pupil time series for subsequent correlation with the BOLD time series. Next, at each temporal window where significant clusters were identified, we (i) transformed standard (MNI) space clusters to subject-specific functional (EPI) space; (ii) converted BOLD responses to percentage of BOLD signal change by scaling with respect to the mean of the BOLD time series; (iii) extracted ROI-level time series by averaging across voxels within the ROI. This yielded ten ROI-level time series for each subject (i.e., six time series at ROIs exhibiting positive activations from 225 to 375 ms, and four time series at ROIs exhibiting negative activations from 375 to 600 ms).

We computed the correlation between pupillary measures and ROI-based BOLD responses after removing the effects of stimulus presentation and behavioral response from both time series. Specifically, (i) the ten ROI-based BOLD responses were all included as regressors in each GLM; (ii) effects of stimulus and RT were removed via linear regression from target BPD/TPR time series; (iii) effects of stimulus, RT, and standard BPD/TPR were removed via linear regression from ROI-based BOLD responses. Significance level of the parameter estimates were determined by a group level one-sample t-test (*P* < 0.01 or 0.05).

#### ROI-based analysis: LC delineation

In preparation for the second and third ROI-based analyses, we delineated LC by taking advantage of a specific imaging protocol (T1-weighted Turbo Spin Echo (TSE)), as well as a set of predefined LC atlases ^64^.

Recent studies have demonstrated that LC has increased neuromelanin concentration, and can be identified in neuromelanin-sensitive TSE scans at bilateral locations adjacent to the fourth ventricle with the most pronounced signal intensities ^12,64^. Using TSE imaging, Keren et al., 2009 ^64^ characterized the spatial distribution of LC and provided two sets of standard space LC templates. Estimated across 44 subjects, these two templates captured the mean, one and two standard deviations (SD) of voxel locations with peak intensity in each axial slice (i.e., 1SD and 2SD LC templates, respectively), therefore providing anatomical references for the location of LC in healthy populations.

In light of these findings, we developed an automated algorithm of LC localization. Specifically, we used the predefined LC atlases to constrain search of voxels with highest signal intensities (i.e., voxels containing LC) in the TSE scans. After registering partial field-of-view TSE scans and standard (MNI) space LC templates to the subject’s structural scans, we compared the average signal intensity in different LC templates to determine voxels which were most likely to contain the LC. We subsequently extracted the time series of LC by computing a weighted average across the two voxels with the highest probability of containing the LC (see Fig. 4 and He et al., 2021 ^85^ for more details). This approach allowed us to extract signal from the “core” instead of the entirety of the LC, with the core of the LC potentially more confined in anatomical and functional heterogeneity ^21,64^.

#### ROI-based analysis: quantifying task-evoked LC responses with EEG

In this analysis, we examined the relationship between temporally specific EEG components and LC activity. At the three temporal windows (225, 275, and 425 ms) where target pupillary measures were significantly correlated with BOLD at ROIs identified with EEG-informed fMRI analysis, BOLD time series at LC were modeled with the same set of five regressors as described in the ***Whole-brain voxel-wise analysis: EEG-informed fMRI analysis***. This analysis was also performed on two additional windows in the temporal vicinity of 225 and 275 ms (i.e., at 250 and 300 ms). Significance level of the parameter estimates were determined by a group level one-sample t-test (*P* < 0.01 or 0.05).

#### ROI-based analysis: quantifying task-evoked LC responses with pupil

In the third analysis, we evaluated the hypothesis that pupil reflects LC activity. We tested this hypothesis on both stimulus type-dependent and stimulus type-independent regressors. In both cases, the GLM included four types of regressors which were convolved with the hemodynamic response function and correlated against the BOLD response at LC: (i) unmodulated stimulus-locked regressors (duration = 0.1 s, height = 1); (ii) duration-modulated RT regressor (onset = target stimulus onset, duration = RT, height = 1); (iii) amplitude-modulated BPD regressor (onset = stimulus onset, duration = 0.1 s, height = BPD of trials); and (iv) amplitude-modulated TPR regressor (onset = stimulus onset, duration = 0.1 s, height = TPR of trials). Significance level of the parameter estimates were determined by a group level one-sample t-test (*P* < 0.01 or 0.05).

It is worth noting that inferences of all analyses in this study were based on results of the target trials (i.e., task-relevant target condition), except when modeling functional responses of LC with pupil. While the relationship between stimulus type-independent BPD (i.e., BPD of both standard and target trials) and BOLD activity at LC was mainly contributed by BPD of standard trials, BPD of target trials exhibited a similar trend with LC activity (although this contribution was not statistically significant when evaluated independently from that of the standard trials, as shown in Fig. 5b). This observation likely reflects the underpowered nature of target trials: with their probability of occurrence one fourth of the standard trials.

#### Data and code availability

Code and data reported in this paper will be made publicly available upon publication.

